# Dimensionality reduction by sparse orthogonal projection with applications to miRNA expression analysis and cancer prediction

**DOI:** 10.1101/2021.11.03.467140

**Authors:** James W. Webber, Kevin M. Elias

## Abstract

**Background:** High dimensionality, i.e. *p > n*, is an inherent feature of machine learning. Fitting a classification model directly to *p*-dimensional data risks overfitting and a reduction in accuracy. Thus, dimensionality reduction is necessary to address overfitting and high dimensionality.

**Results:** We present a novel dimensionality reduction method which uses sparse, orthogonal projections to discover linear separations in reduced dimension space. The technique is applied to miRNA expression analysis and cancer prediction. We use least squares fitting and orthogonality constraints to find a set of orthogonal directions which are highly correlated to the class labels. We also enforce *L*^1^ norm sparsity penalties, to prevent overfitting and remove the uninformative features from the model. Our method is shown to offer a highly competitive classification performance on synthetic examples and real miRNA expression data when compared to similar methods from the literature which use sparsity ideas and orthogonal projections.

**Discussion:** A novel technique is introduced here, which uses sparse, orthogonal projections for dimensionality reduction. The approach is shown to be highly effective in reducing the dimension of miRNA expression data. The application of focus in this article is miRNA expression analysis and cancer predction. The technique may be generalizable, however, to other high dimensionality datasets.

## 1. Background

Often, in machine learning, the data has high dimension, and the dimension is greater than the number of samples, i.e., *p > n*. Fitting a classification model directly to the *p*-dimensional data can lead to overfitting and a reduction in accuracy. The goal in dimensionality reduction is to find a *k*-dimensional subset of the data such that the predictive qualities of the data are retained, without overfitting. In this paper, we consider dimensionality reduction via projections onto *k*-dimensional hyperplanes (i.e., dimensionality reduction by orthogonal projection). For example, consider the synthetic data displayed in figure 1. Here we see a clear linear separation in cases and controls. As an exercise in dimension reduction, we aim to find a one-dimensional subset of the data which retains the linear separability. To do this we consider projections of the data onto one-dimensional planes {*α***w** : *α* ∈ ℝ} (or straight lines, in this case), for some vector **w**. Typically, in early cancer prediction and miRNA expression analysis, a subset of *k* “biomarkers” (or features) would be chosen so as to maximize the separation in cases and controls [19, 16], e.g., using a t-test [9], which can yield poor separation, as is the case in figure 1. In this case, the projection planes are spanned by either **w** = (1, 0) or **w** = (0, 1), and both features, namely miRNA1 and miRNA2, have an *R* = .10 correlation with the class labels.

**Figure 1.**
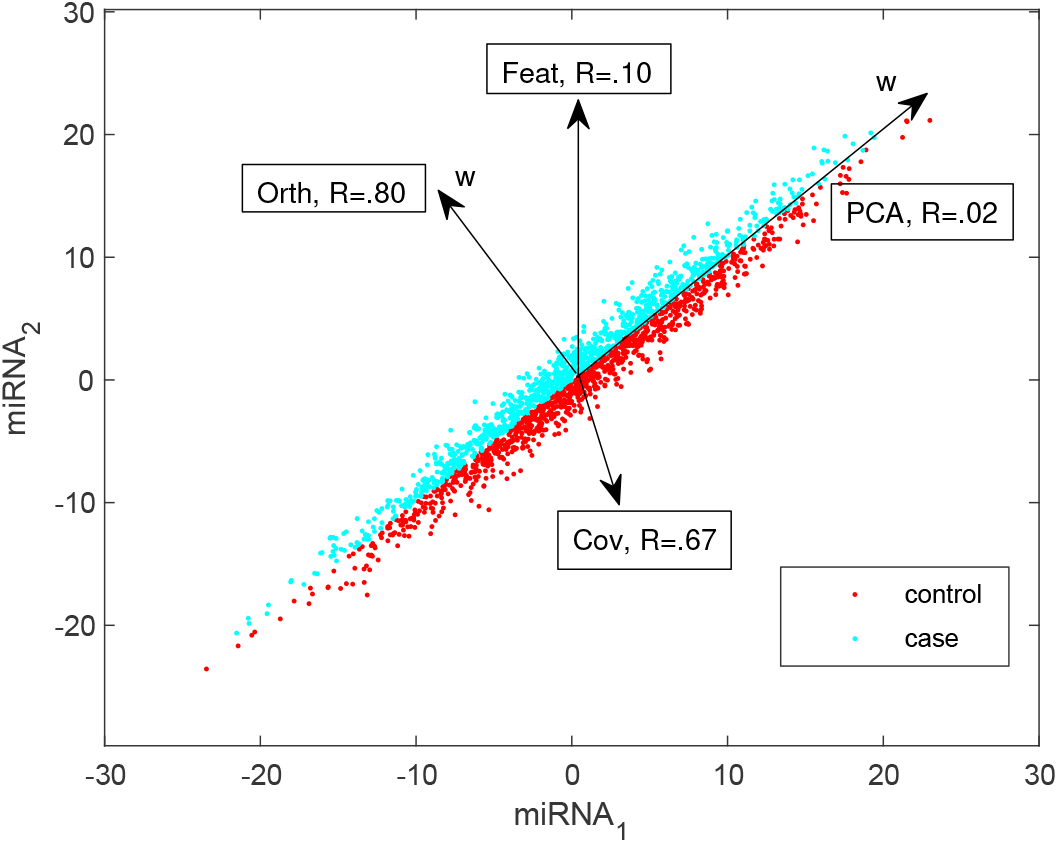
Synthetic data showing possible search directions (**w**) to separate cases and controls. The correlation (R) of the search direction with the controls/case labels is given in each case.

A commonly applied dimensionality reduction method is Principle Component Analysis (PCA) [16, 4], which finds the orthogonal projection of the data which best preserves the variance. Sparse PCA (SPCA) [21], is a generalization of conventional PCA which uses sparsity penalties to select the the most significant features. In the case of figure 1, the direction which best preserves the variance (**w** = (1, 1), denoted “PCA”) is orthogonal to the cancer/control separation, and thus there are cases when the principle components fail to retain the class separation. The literature also considers the Partial Least Squares (PLS) algorithm for dimensionality reduction [18, 11]. The goal of PLS is to find a **w**_1_ such that the covariance between the projected feature and the labels is maximized. Then additional **w**_*i*_, for 2≤ *i*≤ *k*, are computed so that the corresponding extracted features are orthogonal and have high covariance with the labels. In the case of figure 1, the **w** which maximizes the covariance between its projected feature and the labels is denoted “Cov”, which offers an improved class label correlation when compared to feature selection or PCA. Since the PLS directions (**w**_1_, **w**_2_, …, **w**_*k*_) are obtained by simple transpose operations [18], the PLS algorithm is insensitive to overfitting, but at the cost of accuracy. For example, in figure 1, PLS does not find the optimal **w** to separate cases and controls.

In [20], a PLS variant is considered, which uses an orthogonal projection matrix extracted from the PLS algorithm to reduce the data dimension. In conventional PLS, the extracted features are constrained to be orthogonal (as discussed in the last paragraph). However, the matrix which performs the dimensionality reduction is not orthogonal. In [20], the authors argue that, for the purpose of classification, it is most optimal for the dimension reduction matrix to be orthogonal, rather than having orthogonal features. The results of [20] verify the authors hypothesis, and show accurate results when classifying cancers (e.g., leukemia and colorectal cancer) from controls using expression data.

In [8], the authors introduce “DROP-D”, a dimensionality reduction method which is designed for the discrimination of highly correlated, high dimensional (or large *p*) data. DROP-D reduces the dimension by orthogonal projection onto *k*-dimensional hyperplanes, with the goal to preserve the class separation in the reduced dimension space. The class separability is measured in terms of the within and between class dispersions, as is, e.g., done in Fisher Discriminant Analysis (FDA) [5]. DROP-D first cleans the observation matrix of the unwanted features, then the total dispersion matrix of the cleaned data is computed. In the final step, the principle components of the dispersion matrix are used to perform the dimension reduction. In the results presented in [8] on hyperspectral image data, DROP-D offers a similar level of performance when compared to conventional methods, such as PCA and PLS.

The **w** which maximizes the correlation between the projected features and the labels is obtained using least squares fitting. For example, in figure 1, the optimal **w** is labeled “Orth”. In this paper, we use constrained least squares fits to find a set of sparse, orthogonal search directions **w**_1_, **w**_2_, …, **w**_*k*_ with high correlation to the class labels. Following this, we project the data onto the *k*-dimensional hyperplane spanned by **w**_1_, **w**_2_, …, **w**_*k*_ to reduce the dimension. The desired application of this work is early cancer prediction using miRNA expression data. The method we propose is not exclusive to such applications, however, and is directly applicable, in its current form, to any dimensionality reduction problem. To illustrate the technique, we first test the proposed method on synthetic data and real miRNA expression data, then give a comparison to similar methods from the literature. We show that our method offers the most consistent performance in terms of sensitivity and AUC, when compared to the methods of the literature, particularly in the case *p > n*, and when the samples are weighted towards one class (e.g., when there are more controls than cases in the training set). Finally, in section 3, we review implications for our findings and suggest future endeavors.

## 2. Results

The method proposed here will be denoted by Sparse Orthogonal Projection (SOP), for the remainder of this paper. The SOP algorithm and the core objective functions are discussed in detail in the appendix, section A.1. In this section, we present a comparison of SOP, and the methods OPLS [20], FR [3], EFC [13], SPCA [21], and DROP-D [8]. The methods from the literature are discussed in more detail in section A.2. The considered methods are tested on synthetic data, designed to illustrate the effectiveness of SOP, and on real miRNA expression data. The classification model, selection of hyperparameters, and classification metrics used are detailed in the appendix, section A.

### 2.1. A synthetic example; spin top data

Before moving onto real data testing, we introduce some synthetic data here, to provide further motivation for SOP, and which highlights the advantages of SOP when compared to the methods from the literature discussed in section A.2. Consider the data shown in figure 2a. Here we have arranged a 3-D point cloud into the form of a spin top. For more details on the generation of the spin top data, see section A.4. An ideal 2-D projection is shown in figure 2b, which best preserves the class separation and highlights the oscillating boundary. Here we wish to test the effectiveness of SOP and the methods of literature in finding such a 2-D projection as in figure 2b.

**Figure 2.**
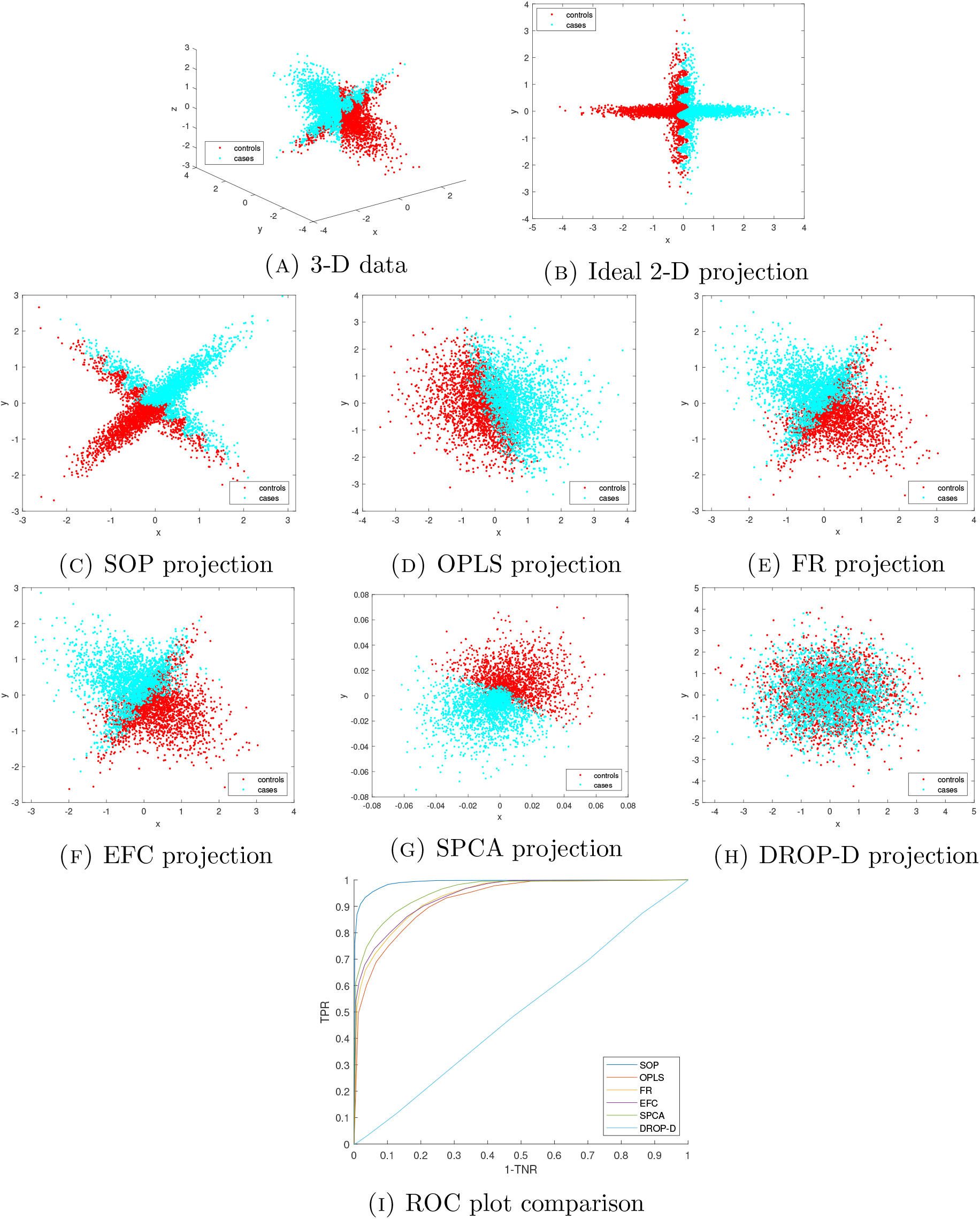
(A) - spin top data. (B) - ideal 2-D projection which preserves the linear separation. (C)-(H) - 2-D projections using the methods considered here for comparison. (I) - ROC plot comparison using each method.

In figures 2c-2h, we have presented the 2-D dimension reductions of the spin top data using SOP, OPLS, FR, EFC, SPCA, and DROP-D. In section B.1, table 2, we present the corresponding classification results, and, in figure 2i, we show ROC plots for each method considered. See section A.4 for details on the classification procedure. In figure 2h, we show the 2-D projection obtained using DROP-D. In this case, DROP-D fails to separate the cases and controls yielding poor AUC and accuracy. This is due to the inability of DROP-D, in this example, to remove the noisy variables. SPCA performs much better and is successful in removing the noisy dimensions. See figure 2g. However, SPCA projects onto the plane perpendicular to the axis of revolution of the spin top, which merges some cases and controls along the oscillating boundary. In figures 2e and 2f we see that the feature selection methods, FR and EFC, successfully remove the noisy variables, and offer the same 2-D projection (i.e., the same set of two features are selected using FR and EFC). However, the spin top is misoriented in the 2-D projection, which yields some overlap in cases and controls, and thus a reduction in classification accuracy. In figure 2d, the OPLS projection is corrupted by the noise, which causes a significant overlap in cases and controls. We see only mild noise effects and misorientation of the spin top in figure 2c using SOP, which outperforms the methods from the literature across all metrics considered.

### 2.2. Results - Japanese data

Here we present our results on the miRNA expression data of [19, 16, 14], collected from Japanese patients. This data is discussed in point of section A.5. For each of the 15 diseases considered in [19, 16, 14], we perform a separate, binary classification, where we aim to separate the given disease from the control set.

See figures 3a-3d, where we have shown box plots of our classification results over all 15 diseases, and table 1 where we report the mean and standard deviation classification scores. The results in table 1 are presented in the form *µ*± *σ*, where *µ* is the mean score across all 15 diseases and *σ* is the standard deviation.

**Figure 3.**
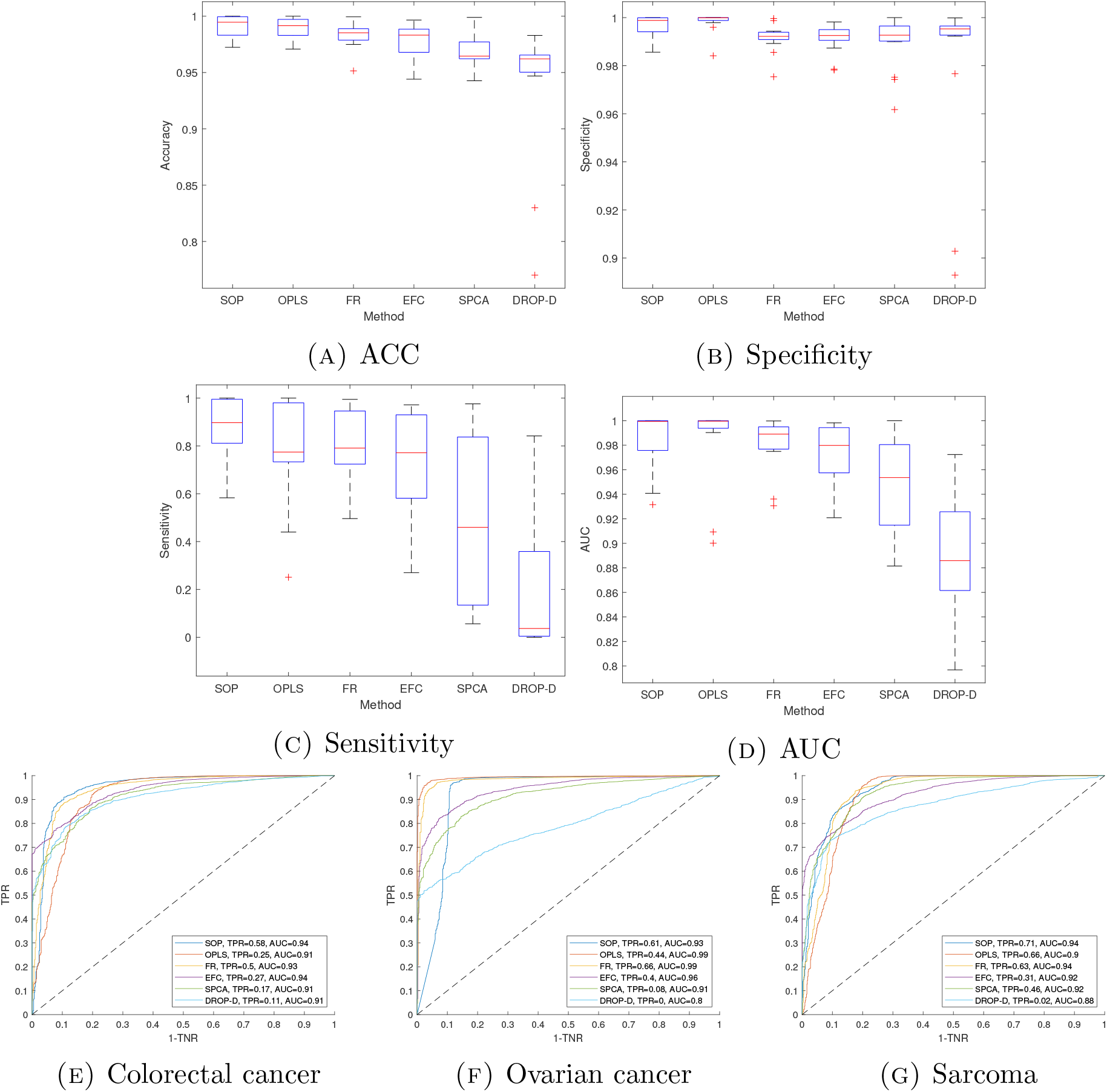
(A)-(D) - Box plots showing the median, upper and lower quartiles, and minima and maxima, across 15 classification results on the Japanese data set, where each of the 15 diseases was separated from the control set. The outliers are highlighted as red crosses. OPLS and FR have two AUC outliers corresponding to the colorectal cancer and sarcoma classifications. SOP has one AUC outlier corresponding to the ovarian cancer classification. (E)-(G) - ROC comparisons corresponding to each AUC outlier. The sensitivity scores and AUC values for each method are given in the figure legend in (E)-(G).

**Table 1.** Japan results. In the table, ∼1 and ∼0 indicate that the score is one or zero, respectively, to two significant figures.

| Method | AUC | $F_1$ | ACC | TNR | TPR |
| --- | --- | --- | --- | --- | --- |
| SOP | $.98 \pm .03$ | $.89 \pm .12$ | $.99 \pm .01$ | $\sim 1 \pm \sim 0$ | $.87 \pm .14$ |
| OPLS | $.99 \pm .03$ | $.85 \pm .17$ | $.99 \pm .01$ | $\sim 1 \pm \sim 0$ | $.78 \pm .21$ |
| FR | $.98 \pm .02$ | $.82 \pm .12$ | $.98 \pm .01$ | $.99 \pm .01$ | $.81 \pm .15$ |
| EFC | $.97 \pm .02$ | $.75 \pm .20$ | $.98 \pm .02$ | $.99 \pm .01$ | $.72 \pm .24$ |
| SPCA | $.95 \pm .04$ | $.52 \pm .34$ | $.97 \pm .02$ | $.99 \pm .01$ | $.46 \pm .36$ |
| DROP-D | $.89 \pm .05$ | $.25 \pm .30$ | $.94 \pm .06$ | $.98 \pm .03$ | $.21 \pm .29$ |

**Table 2.** Spin top data results.

| Method | AUC | $F_1$ | ACC | TNR | TPR |
| --- | --- | --- | --- | --- | --- |
| SOP | .99 | .95 | .95 | .95 | .95 |
| OPLS | .92 | .84 | .84 | .86 | .81 |
| FR | .94 | .84 | .85 | .85 | .84 |
| EFC | .94 | .85 | .85 | .86 | .84 |
| SPCA | .96 | .88 | .88 | .88 | .88 |
| DROP-D | .50 | .52 | .50 | .48 | .52 |

In this example, SOP, OPLS, and FR perform well in terms of the classification accuracy, AUC and specificity, with SOP offering the best performance in terms of mean sensitivity and *F*_1_ score. The standard deviation *F*_1_ scores and sensitivities offered by SOP, are less than or equal to those of FR and OPLS, indicating that SOP offers a more consistent performance in terms of *F*_1_ score and sensitivity. The box plots of figure 3 also support this. In particular, OPLS has an outlier sensitivity score of TPR = .25, which corresponds to the colorectal cancer classification. FR offers a significantly higher sensitivity of TPR = .50 on the colorectal cancer classification, with SOP offering the best sensitivity of TPR = .58. SOP has a longer lower AUC tail in the box plots, when compared to OPLS and FR, which is an advantage for OPLS and FR over SOP. In particular, in the ovarian cancer classification, SOP offers an AUC of AUC = .93. whereas OPLS and FR both yield AUC’s of AUC = .99. See figure 3f. OPLS and FR share two AUC outliers, corresponding to the colorectal cancer and sarcoma classifications. In figures 3e and 3g, we present ROC comparisons for the colorectal cancer and sarcoma classifications. The two outlier AUC’s for OPLS are lower than the SOP outlier. The minimum AUC offered by OPLS is AUC = .90, for the sarcoma classification, whereas the minimum AUC for SOP is AUC = 93 for ovarian cancer. This is an advantage for SOP when compared to OPLS. The FR AUC outliers are similar to those of SOP.

We see a reduced classification performance using EFC, SPCA, and DROP-D, when compared to SOP, OPLS, and FR. In particular, the mean sensitivity and *F*_1_ score is low for EFC, SPCA, and DROP-D. For the majority (13 out of 15) of the classifications performed here, the number of cases was significantly smaller than the control set. For example, there are 119 breast cancer patients in total with 1123 controls. As noted in the scatter plots of the Japanese data discussed in appendix C, a large proportion of the control set is well separated from the cancers, and thus a high specificity can be achieved using all methods. SOP is more effective in handling the mismatch in class sizes, and offers a higher mean sensitivity and *F*_1_ score, with lower standard deviation, when compared to the similar methods from the literature.

### 2.3. Results - Lee *et. al*. data

In this section we present our results on the data provided by Lee *et. al*. in [12], i.e., the data discussed in point (3) of section A.5. In figure 4, we present an ROC plot comparison using SOP, OPLS, FR, EFC, SPCA, and DROP-D. In section B.1, table 3, we report the classification scores discussed in section A.3.

**Figure 4.**
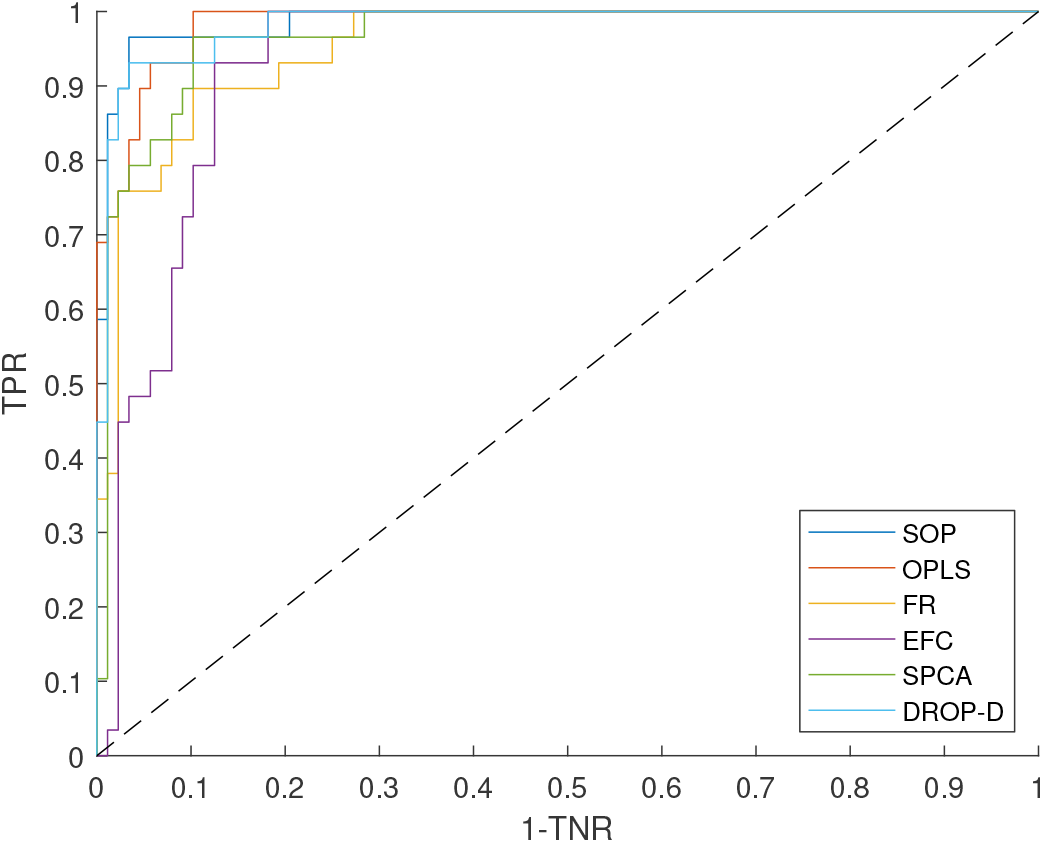
ROC plot comparison on Lee *et. al*. data.

**Table 3.**
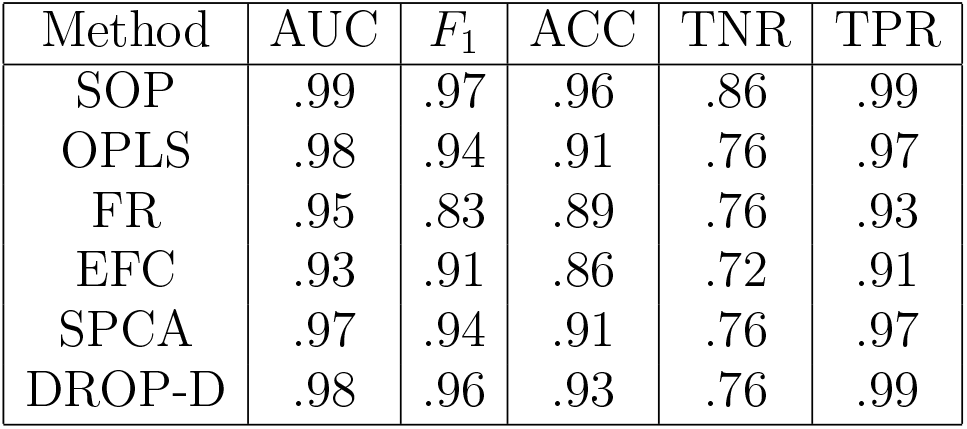
*Lee et. al*. data results.

| Method | AUC | $F_1$ | ACC | TNR | TPR |
| --- | --- | --- | --- | --- | --- |
| SOP | .99 | .97 | .96 | .86 | .99 |
| OPLS | .98 | .94 | .91 | .76 | .97 |
| FR | .95 | .83 | .89 | .76 | .93 |
| EFC | .93 | .91 | .86 | .72 | .91 |
| SPCA | .97 | .94 | .91 | .76 | .97 |
| DROP-D | .98 | .96 | .93 | .76 | .99 |

In this example all methods perform well, with SOP yielding the best results. In particular, SOP offers a specificity of TNR = .86, a 10% increase over the next best performing method. In terms of specificity, the second best performing methods are OPLS, FR, SPCA, or DROP-D, which offer a specificity of TNR = .76. SOP also offers greater than or equal AUC, *F*_1_, ACC, and TPR scores, although the difference is more minor for these statistics. For example, SOP offers a classification accuracy of ACC = .96, a 3% increase over the next best performing method, namely DROP-D, which offers an accuracy of ACC = .93. As was often the case with the Japanese data classifications conducted in section 2.2, the number of cases and controls in the Lee *et. al*. data is significantly different (29 controls and 88 cases). Although in this example there are more cases than controls, whereas the Japanese data was weighted towards controls. In section 2.2, it was harder to achieve high sensitivity given the weighting towards controls. In this case, the specificity is of concern given the weighting towards cases. This example provides further evidence that SOP is most optimal for data sets weighted to one class, when compared to the methods of the literature considered here.

### 2.4. Multi-classification results

In this section we consider the case when *n*_*c*_ *>* 2. We aim to test the effectiveness of SOP, and the methods of the literature, in separating controls from breast cancer, and advanced breast cancer patients simultaneously in a multi-classification. In this case *n*_*c*_ = 3. The data used here is extracted from the miRNA expression data provided by the authors in [14, 19, 16]. For more details, see section A.6.

See figure 5 where we have plotted the ROC curves for each method considered. Here we plot AUC curves for each class separately, for each method, to give a more detailed breakdown of the performance of each method. In section B.1, table 4, we have presented the mean classification scores over all classes, for each method considered.

**Figure 5.**
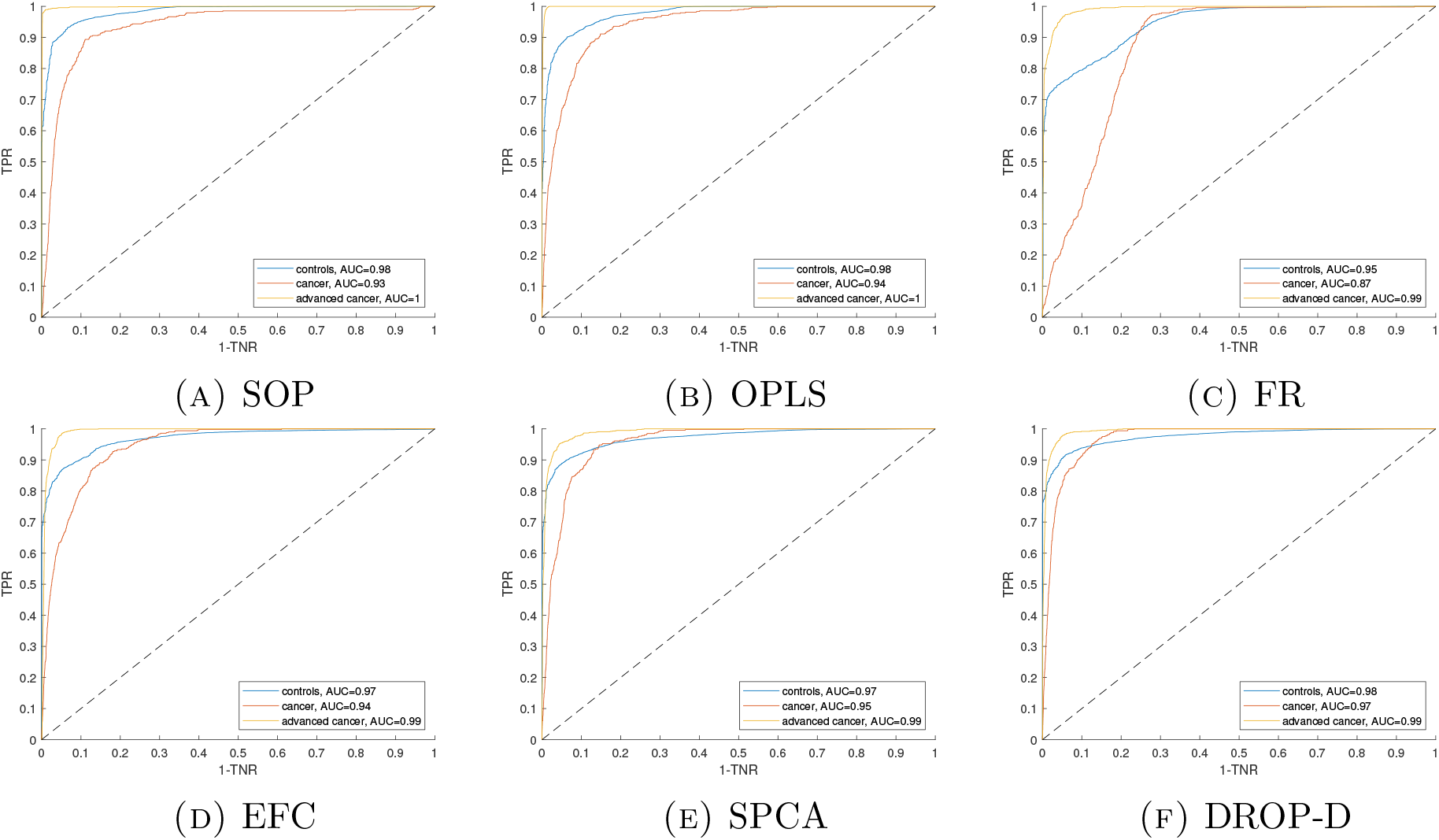
ROC plots for the breast cancer multi classification, separating controls from cancer and advanced cancer. The AUC scores corresponding to each class are given the figure legends to two significant figures.

**Table 4.** Japan breast cancer multi-class results. The scores presented are the mean over all *n*_*c*_ = 3 classes.

| Method | AUC | $F_1$ | ACC | TNR | TPR |
| --- | --- | --- | --- | --- | --- |
| SOP | .97 | .78 | .97 | .93 | .79 |
| OPLS | .97 | .77 | .96 | .94 | .84 |
| FR | .93 | .62 | .97 | .86 | .61 |
| EFC | .97 | .69 | .95 | .87 | .67 |
| SPCA | .97 | .72 | .96 | .88 | .70 |
| DROP-D | .98 | .73 | .97 | .88 | .70 |

In this example, OPLS and SOP offer the best overall performance across all metrics. DROP-D yields the best AUC score. In particular, DROP-D offers the optimal AUC score (AUC = .97) on the cancer patient class. See the red ROC curves of figure 5. OPLS offers the highest mean sensitivity of TPR = .84, and SOP offers the highest *F*_1_ score of *F*_1_ = .78. We see a reduced performance in terms of mean *F*_1_ score, specificity and sensitivity using FR, EFC, SPCA, and DROP-D, when compared to SOP and OPLS. This example highlights the effectiveness of SOP, and other similar methods, such as OPLS, in classifying patients into *n*_*c*_ *>* 2 groups.

## 3. Discussion

This study introduced the dimension reduction methodology, SOP. In section 2, we tested the effectiveness of SOP, and the methods from the literature, on synthetic data, and real miRNA expression data publicly available in the literature [19, 16, 14, 12], with the goal to distinguish control patients from those with cancer. We introduced a novel, synthetic “spin top” data set in section 2.1, designed for testing dimensionality reduction methods which use orthogonal projections. SOP was shown to produce optimal results on the spin top data set, across all metrics considered. In the examples conducted on real miRNA expression data, SOP offered a competitive or greater performance than the methods of the literature, particularly in cases when the class sizes were unbalanced (e.g., if more controls than cases were available in the training set), and when the data dimension (*p*) was high. In such situations, SOP was most effective in correctly classifying patients into the class with fewer training samples. In total, we considered three different *p* values, namely *p* = 103 (spin top data), *p* = 2565 (Japanese data), *p* = 2578 (Lee *et. al*. data). In sections B.2.1 and B.2.2, we present results on two more real miRNA expression data sets from the literature, namely that of Keller *et. al*. [9] *(in this case p* = 863), and Chan *et. al*. [2] *(p* = 280). These results are included to test the performance of SOP more rigorously across a range of (*n, p*) values. The choice for the sparsity parameter *β* introduced here (see section A.9) appears to be a good one, as evidenced by the results. SOP, with *β* chosen as in section A.9, performs well in all examples considered, and across a variety of (*n, p*). In section 2, the number of features (*k*) was chosen to give the best result in terms of mean AUC across all classifications considered in each case. In the appendix, section B.3, we address the tuning of *k*, and perform the same set of classifications as in section 2, using SOP, but with *k* = 3 fixed throughout all classifications.

Typically, in cancer prediction using miRNA expression, binary classification problems are considered [20, 19, 16, 14, 12, 2], and methods such as OPLS and EFC are applied to separate the two classes. While the majority of the classifications conducted in this paper were binary (i.e., the number of classes *n*_*c*_ = 2), in section 2.4 we also tested the effectiveness of SOP on a multi-classification problem, where *n*_*c*_ = 3, and the goal was to separate control patients from those with breast cancer and advanced breast cancer simultaneously. In this example, SOP offered a competitive classification performance when compared to the best approach, OPLS. These results provide evidence that such dimension reduction methodologies can be applied effectively to multiple classification problems, i.e., when the number of classes *n*_*c*_ *>* 2.

## 4. Conclusions and further work

In this paper we introduced SOP, a novel algorithm for dimensionality reduction using sparse orthogonal projections. The desired application of this work is miRNA expression analysis and cancer prediction. SOP was compared against five similar methods from the literature which use orthogonal projections and sparsity constraints, namely, OPLS [20], FR [3], EFC [13], SPCA [21], and DROP-D [8]. The technique was proven to be highly effective in reducing the dimension of miRNA expression data, and offered a highly competitive performance when compared to the methods of the literature. The technique may be applicable to other high dimension data sets (e.g., gene expression arrays). Further analysis is needed, however, to validate the performance on other high dimension data sets.

In further work we aim to test the effectiveness of SOP in the case when *n*_*c*_ *>* 3, as only *n*_*c*_ = 2, 3 are considered here. We suspect that the classification accuracy will decrease with increasing *n*_*c*_, as it is more difficult to find linear separability for multiple classes simultaneously. Further testing is needed to confirm this hypothesis. In many of the examples conducted in this paper, we assumed knowledge of a high number of miRNA expressions in order to fit our classification models. For example, *p* = 2565 out of a possible *p* = 2585 [10, chapter 3] miRNA’s were used to classify Japanese patients with cancer and liver diseases (e.g., hepatitis) in section 2.2. With such rich data available, we saw a high classification performance, particularly when using OPLS, FR and SOP. In practice, however, it may not be possible to measure all miRNA expression values for every patient. Thus, in further work, we aim to consider the case when not all expression values are known, and what can be done here to retain the classification performance. Specifically, we aim to consider a data imputation to fill in the missing values in situations where we have limited data available.

## 5. Declarations

Here we give our declarations.

### 5.1. Ethics approval and consent to participate

No new human or animal data is presented here.

### 5.2. Consent for publication

There are no issues regarding consent for publication.

### 5.3. Availability of data and materials

All real miRNA expression data sets considered here are publicly available online. See section A.5 for more details. The SOP code and spin top data introduced here is available from the authors upon reasonable request.

### 5.4. Competing interests

There are no financial or non-financial competing interests.

### 5.5. Funding

This research received support from the grant K12 HD000849, awarded to the Reproductive Scientist Development Program by the Eunice Kennedy Shriver National Institute of Child Health and Human Development (KME). The authors also wish to acknowledge funding support from the GOG Foundation, as part of the Reproductive Scientist Development Program (KME), Robert and Deborah First Family Fund (KME), the Massachusetts Life Sciences Center Bits to Bytes Program (JWW, KME), and Abcam, Inc (JWW).

### 5.6. Authors’ contributions

JWW developed SOP and the original idea. The experiments were conducted by JWW. Analyses of results by JWW and KME. KME was as major contributor in writing the manuscript, and provided expert insight from a medical background needed to communicate this work to a medical audience.

## Appendix A

### Methods

In this section we introduce our dimensionality reduction strategy, SOP, and discuss the data sets that will be used for testing. We also discuss the methods from the literature that we compare our method against, and introduce a new, synthetic data set to test the performance of the proposed method, and those of the literature.

#### A.1. Dimensionality reduction methodology

Throughout this paper, we let *X* = (**x**_1_, …, **x**_*p*_) ∈ℝ^*n×p*^ denote a matrix of input variables (e.g., miRNA expression data), and **y** ∈ **{**0, …, *n*_*c−*_ 1} ^*n*^ a vector of class labels, where *n*_*c*_≥ 2 is the number of classes, *p* is the dimension of the data (i.e., the number variables), and *n* is the number of samples. We let *k < p* denote the number of features extracted after a dimensionality reduction.

We aim to perform dimensionality reduction by right multiplication of *X* with a sparse, orthogonal matrix *W* ∈ ℝ^*p×k*^, and extract a set of features *T* ∈ ℝ^*n×k*^ via

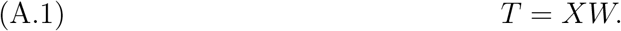

The calculation of *W* can be broken down into two steps. First, we find

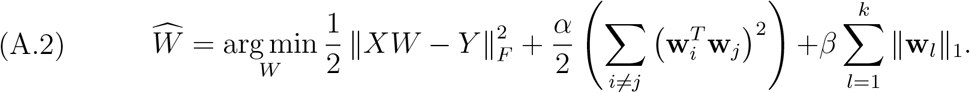

where 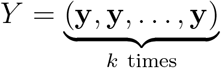 and ‖ · ‖·_*F*_ is the Frobenius norm. Then we decompose 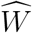 as its Singular Value Decomposition (SVD)

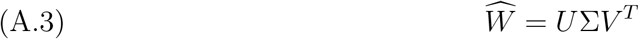

and set *W* = (**u**_1_, …, **u**_*k*_)(**v**_1_, …, **v**_*k*_)^*T*^. That is, we first find a sparse, orthogonal matrix 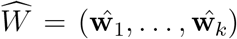, for which 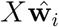, for every 1 ≤ *i* ≤ *k*, has high correlation with the labels **y**. Then, we set *W* as the closest orthogonal matrix to 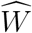 (in terms of Frobenius norm) in the second step using the SVD of 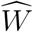. After *W* is calculated, and the features *T* = *XW* are extracted, a classification model (e.g., softmax function, support vector machine) can be fit to the reduced dimension data. The first term of A.2 fits the input variables *X* to the response *Y*, and finds **w**_*i*_ for which *X***w**_*i*_ has high correlation with **y**. The *α* term is included to ensure orthogonality of *W*. In practice, we set *α* to some large value, orders of magnitude greater than the maximum expression, so that *W* is hard constrained to be orthogonal. The *β* term is included to enforce sparsity and prevent overfitting. This follows the idea that many of the variables will have little predictive quality, which is typically the case in miRNA expression analysis and cancer prediction [4]. Hence, the *L*^1^ penalty is included to remove the noisy, less informative variables from consideration in the model.

We now give specific details on the implementation of SOP. To solve the objective in equation (A.2) and calculate 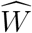, we apply the L-BFGS-B code of [1]. L-BFGS-B requires an objective function and its gradient. All elements of equation (A.2) are smoothly differentiable with the exception of the *L*^1^ norm, and the gradients can be calculated easily for L-BFGS-B implementation. To calculate the gradient of the *L*^1^ norms, we use the subgradient of the absolute value function

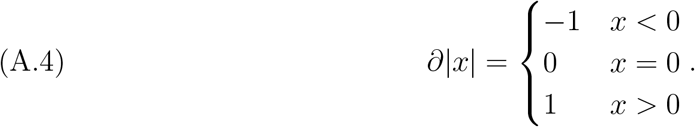

To initialize L-BFGS-B we use the classical solution to the orthogonal procrustes problem [7]. Specifically, we implement the algorithm, SOP, below:

1. Set the input matrix *X*, the label matrix *Y*, and the number of features to be extracted *k*. Set *α* and *β*.
2. Center and normalize the input data *X*. For example, in section 2, the data was normalized to the cube [−1, 1]^*p*^, and centered with zero mean.
3. Find

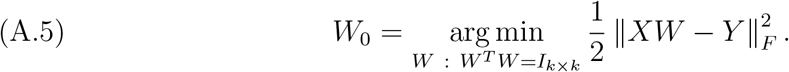

Let *M* = *Y* ^*T*^ *X*, and let *M* = *U* Σ*V* ^*T*^ be decomposed as its SVD. Then, *W*_0_ = (**v**_1_, …, **v**_*k*_)(**u**_1_, …, **u**_*k*_)^*T*^ [7].
4. *Reassign* 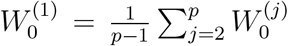, where 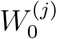 is row *j* of *W*_0_. We do this reassignment as the entries of 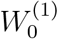 were typically much higher magnitude than the remaining row entries in the procrustes solution. Hence we reassign 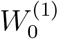 to the average row to give more equal footing to each of the *p* features at the initialization stage.
5. Solve the objective A.2 using L-BFGS-B, with the normalized *X*, and the label matrix *Y* as input, setting *W*_0_ as an initial guess, *α* as the orthogonality constraint parameter, *β* as the sparsity parameter, and *k* as the number of features. Output 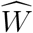.
6. Decompose 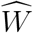 into its SVD, 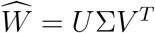, and output *W* = (**u**_1_, …, **u**_*k*_)(**v**_1_, …, **v**_*k*_)^*T*^. The SOP code detailed above is available from the authors upon request.

#### A.2. Methods for comparison

Here we discuss the methods from the literature which we compare the proposed method (SOP) against. We choose to compare against other methods which use sparsity ideas and orthogonal projections, with the idea to compare against other methods which are most similar to SOP.

We compare against the PLS variant of [20], since the idea to use orthogonal projections which retain linear separations bears similarities to SOP. We denote this method as Orthogonal PLS (OPLS). DROP-D searches for linear separability and uses orthogonal projections, similarly to SOP, and thus why it is a chosen point of comparison. We compare against Sparse PCA (SPCA) [21], a generalization of conventional PCA which uses sparsity penalties to select the the most significant features. To implement SPCA in the comparisons we apply the code of [15]. We also compare against two commonly applied feature selection methods, namely Forward Regression (FR) [3], and Expression Fold Change (EFC) [13]. FR is similar to a correlation based feature selection, commonly applied in cancer prediction [4], except in FR the features are chosen to have low linear dependence with one another, which is not a consideration in oblivious feature selection. EFC computes the ratios between the mean case and control sample. Then the features with mean ratio farthest from 1 are ranked most highly. EFC is a commonly applied idea in miRNA expression analysis and cancer prediction [13].

#### A.3. Classification metrics

Here we introduce the metrics which will be used to assess the quality of the results. Let TP, FP, TN, and FN denote the number of true positives, false positives, true negatives, and false negatives, respectively, in a binary classification. Then, we report the following classification metrics in our comparisons:

1. The classification accuracy (denoted by “ACC”)

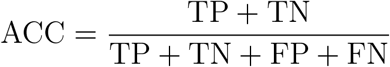
2. The *F*_1_ score

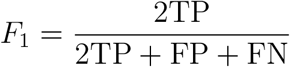
3. The True Positive Rate (TPR, also known as sensitivity)

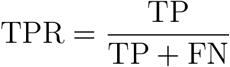
4. The True Negative Rate (TNR, also known as specificity)

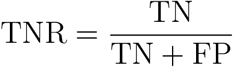

We also show plots of Receiver Operator Characteristic (ROC) curves, and report the Area Under the Curve (AUC) values. All metrics reported here take values between 0 and 1. A value closer to 1 indicates a better performance, and vice versa.

#### A.4. Spin top data

Here we discuss in more detail the spin top data introduced in section 2.1, and the classification methods used in the results presented. The spin top data is formed by merging two Gaussian point clouds, one with low standard deviation in the *z* axis direction (i.e., the flat plane perpendicular to the axis of rotation of the spin top), and one with low standard deviation in the (*x, y*) plane (i.e., the thin tube parallel to the axis of rotation). The spin top is then rotated about the origin to form the point cloud shown in figure 2a. There are two classes, which are separated by an oscillating sinusoid boundary. To make the dimensionality reduction more challenging, we embed the 3-D spin top data into 103-dimensional space by adding 100 dimensions of Gaussian noise. So, in total, there are *p* = 103 dimensions, only 3 of which have any relation to the control/case separation, and *n* = 4500 samples. The goal of the spin top problem is to remove the noisy variables, while simultaneously projecting the spin top onto a 2-D plane (e.g., as in figure 2b) so that the classes separate.

To clarify the classification process in section 2.1, for each method the number of features was set to *k* = 2. After the dimension reduction was performed, we fit a *K* nearest neighbors classifier with *K* = 11 neighbors. Then, we generated a test data set (size *n* = 4500) from the spin top distribution of figure 2a, input the test data into our model, and calculated the classification metrics discussed in section A.3 for each method. For SOP, we set *α* = 10^6^ and *β* = 10^2^ (see equation A.2). For SPCA we set *λ* = *λ*_1,*j*_ = *λ*_1,*i*_ (the sparsity parameter) for every 1 ≤ *i,j* ≤ *k* (using the notation of [21]), and chose *λ* to give the best results in terms of AUC. When implementing DROP-D, we set *b* = 2, and chose the *w* (using the notation of [8]) which gave the best results in terms of AUC. We set *b* = 2 as this was shown to produce good results in the examples conducted in [8].

#### A.5. Real data sets

We consider the following four data sets from the literature for real data testing:

1. Serum miRNA expression data of [19, 16, 14] collected from 2460 Japanese patients. Here the authors provide expression values for 1123 control patients, and for patients with 15 different types of diseases (the case number varies with the disease), including bladder cancer, hepatocellular carcinoma (HCC), breast cancer, ovarian cancer, and hepatitis. To compile the data used here, we combine the data sets of [19] and [16], making sure to delete any replicate patients. In [14] expression values are provided for 79 patients with advanced breast cancer. We combine the advanced breast cancer samples of [14] with the breast cancer samples of [19, 16]. In [14], the number of expression values recorded is *p* = 2535, which is slightly smaller than the *p* = 2565 of [19, 16]. For the breast cancer classifications conducted here, we use the *p* = 2535 miRNA’s of [14] so that the data sets can be combined. For the remaining diseases, we use the full set of *p* = 2565 miRNA expression values available, e.g., in our HCC classifications. In appendix C, we give a statistical and visual analysis of the Japanese data, and discuss some of its properties.
2. The expression data of [9] collected by Keller *et. al*. from 454 German patients. The data is comprised of 70 healthy control patients, and patients with 14 different cancer and noncancer diseases (the case number varies with the disease), including lung cancer, ovarian cancer, multiple sclerosis, and pancreatic cancer. The number of recorded expression values, in this case, is *p* = 863, for all patients. One of the cancers studied in [9] were Wilms tumors, for which the authors provided 5 case samples. As the number of case samples for Wilms tumors is low, we do not consider the Wilms tumor cases in our analyses. So, in total, we consider 13 different diseases from the Keller data set (i.e., the 14 provided minus the Wilms tumor samples).
3. miRNA expression data of [12] collected by Lee *et. al*. from 232 Korean patients. This data is comprised of 88 patients with Pancreatic Cancer (PC), and 19 healthy controls. In [12], the authors combine the 19 healthy patients with 10 cholelithiasis patients to form a larger control set of 29 patients for use in the PC classifications. We use the same control set here, and aim to separate the controls from the PC patients. In total we consider *n* = 117 patients. The authors provide *p* = 2578 expression values, for each patient, all of which will be used in our classifications.
4. RT-PCR data of [2] collected by Chan *et. al*. from 54 Singaporean patients. The data is comprised of 32 patients with breast cancer and 22 healthy controls. The authors provide *p* = 280 expression values for each patient. We consider all patient samples and expression values provided, and hence in this case, *n* = 54 and *p* = 280.

#### A.6 Multi-class data

The Japanese breast cancer samples come from three separate sources, namely [14, 19, 16]. In [14], the patients are all diagnosed with advanced breast cancer, whereas in [19, 16] the cancer stage is not specified. In section C.1, we present scatter plots showing two distinct clusters for the cancer patients of [19, 16] and [14]. Given the distinct clustering among cancer patients, and the controls, it is reasonable to consider a multi-classification, whereby we aim to classify patients into *n*_*c*_ = 3 classes simultaneously. In this case, the three classes are controls, breast cancers, and advanced breast cancers.

EFC is only defined when *n*_*c*_ = 2. To remedy this, and generalize EFC for *n*_*c*_ *>* 2, we calculate the EFC value for every possible pair of classes and rank the features based on the mean fold change over each pair. In this example, we calculate the EFC for control vs cancer, control vs advanced cancer, and cancer vs advanced cancer, and take the average EFC. Then, once we have assigned a single fold change value to each feature, the features are ranked in the same way as conventional EFC, and the top *k* features based on ranking are extracted. There may be a more optimal way to adapt EFC to multi-classification. However, such updates to EFC are outside the scope of this paper. For each method considered, we use the optimal *k* ∈{2, …, 10} chosen for the binary classifications conducted on the Japanese data set in section 2.2. The remaining hyperparameters are chosen as discussed in section A.9.

#### A.7. Classification model used for real data testing

Here we discuss the classification model used to classify patients in the real miRNA expression data experiments conducted in this paper. Let *T*∈ℝ^*n×k*^ be a set of *k* features extracted using one of the dimensionality reduction methods considered (e.g., OPLS). To classify patients, we train a softmax function [6] classification model

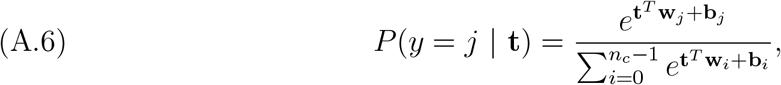

where *j*∈{0, 1, …, *n*_*c*_ *−* 1} is the class label, **t** ℝ^*k*^ is a sample in the reduced dimension space (e.g., one row of *T*), and the (**w**_*j*_, **b**_*j*_) are weights and biases to be trained. Here *y* denotes the class label assigned to **t**. The class with the highest probability *P* is then chosen for membership. A softmax function is typically used as the final layer in a neural network to assign a probability of membership to each class [6]. Typically neural nets have a hierarchical structure, and the number of hidden neurons decreases with each added layer, with the idea to reduce the data dimension before the softmax classification. We use a similar idea here, except the dimensionality reduction is implemented using one of the methods discussed in section A.2, rather than by intermediate hidden network layers. The softmax function classification model is used throughout the real miRNA expression data examples conducted in this paper.

#### A.8. Validation methods used for real data testing

Here we discuss the validation methods used in the real miRNA expression experiments conducted. For the data sets with small *n*, i.e., data sets (2)-(4) of section A.5, where *n <* 150, we use a Leave One Out (LOO) cross validation. For data set (1) of section A.5, where *n >* 1000, we use a multiple hold out set validation. That is, we randomly and uniformly set aside a subset of samples size *n*_*t*_ *< n* from the data, fit a model to the remaining *n* − *n*_*t*_ samples, and validate the performance on the hold out set. Then the process is repeated *n*_*T*_ times and the results are averaged. We set *n*_*t*_ = 300 and *n*_*T*_ = 50, so in total there are *n*_*T*_ × *n*_*t*_ = 15000 test trials for each classification performed on data set (1).

#### A.9. Hyperparameter selection and normalization methods used for real data testing

Throughout the experiments conducted here on real miRNA expression data, all data were normalized to the cube [−1, 1]^*p*^ and centered with mean zero. For SOP, we set *α* = 10^3^ throughout the simulations conducted in this section. *α* = 10^3^ is such that the orthogonality constraint of A.2 is satisfied to 3 significant figures, which is sufficient, as in step two of SOP, discussed in section A.1, the projection matrix is reassigned to the closest orthonormal matrix using the SVD. Thus, the matrix used in the SOP dimensionality reduction is orthonormal to machine precision. In general, we find that setting *α* = 10^*m*^ yields a 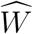 in the first step of SOP, which is orthogonal to *m* significant figures of accuracy, when using the proposed normalization strategy. The sparsity parameter *β* is set to

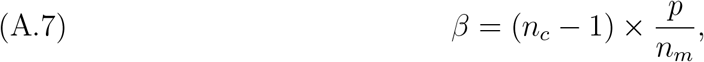

where *p* (the dimension) is the number of available miRNA expression values, *n*_*c*_^≥^ 2 is the number of classes, and *n*_*m*_ = 2585 is the total number of all known miRNA’s in humans [10, chapter 3]. The idea here, is that for larger *p* there will be a higher proportion of noise and uninformative expression values in the data. Thus, we scale *β* according to A.7 based on the number of expression values we have available for training. We multiply by *n*_*c*_ − 1 in (A.7), so that *β*, and the *L*^1^ penalization, scales better with the max value of *Y* (i.e., *n*_*c*_ − 1) and the magnitude of the least squares error term of (A.2). We find that such choice for *β* works well when paired with the proposed normalization strategy, for all examples conducted in this section. For SPCA, we set the sparsity parameter in the same way, for fairness. Also, in the SPCA implementation, soft thresholding is used here to calculate the components, as is recommended in [21] for the case when *p > n*, and for gene expression arrays. All data sets considered in this section satisfy *p > n*.

In the DROP-D implementation, we set *b* = 2, and *w* = 5 (using the notation of [8]), as these values were shown to yield good results on the examples presented in [8]. For all methods, we set the number of features (*k*) to give the best results in terms of AUC on the test set, over all *k*∈{2, 3, …, 10}. For the Keller and Japanese data sets (data sets (1) and (2) of section A.5), we chose the *k* ∈2, 3, …, 10 which gave the best mean AUC across all diseases considered. For example, on the Japanese data, we performed 15 sets of binary classifications with the aim to separate the given disease from the control set, and chose the *k* ∈{2, 3, …, 10} which gave the best mean AUC over all 15 classifications, after cross validation.

## Appendix B

### Additional results

#### B.1. Additional tables

Here we present additional tables for the results of section 2, for each data set considered. The Japanese data results table is not included, since it was already shown in the main text.

#### B.2. Additional binary classification results

Here we present additional comparisons of the considered methods on the miRNA expression data of Keller *et. al*. [9] *and Chan et. al*. [2].

##### B.2.1. Results - Keller et. al. data

Here we present our results on the data provided by Keller *et. al*. in [9]. This data is discussed in point (2) of section A.5. For each of the 13 diseases considered (note that we are excluding Wilms tumors due to the low number of cases), we perform a binary classification, where we aim to separate the given disease from the control set, as was done with the Japanese data considered in section 2.2.

See figure 6, where we have shown box plots of our classification results over all 13 diseases, and table 5 where we report the mean and standard deviation classification scores. As in section 2.2, the results in table 5 are presented in the form *µ* ±*σ*, where *µ* is the mean score across all 13 diseases and *σ* is the standard deviation. The results indicate an improved accuracy and specificity using OPLS when compared to SOP, whereas SOP offers an improved sensitivity and a more consistent AUC. As was the case with the Japanese data, discussed in section 2.2, SOP offers better sensitivity than the methods of the literature. Ultimately, in this case, the optimal classifier is dependent on the situation. For example, we may wish to screen patients for risk of cancer, and decide who should go forward for further testing (e.g., imaging). In this case we do not want to miss any patients who have cancer or may go on to develop cancer (i.e., we want high sensitivity), and SOP would be most appropriate. In other cases, for example if we are recommending that patients have surgery based on the predicted risk of cancer, we would like to avoid admitting patients to surgery if they are not to develop cancer (i.e., high specificity is desired), and OPLS would be the most appropriate model.

**Figure 6.**
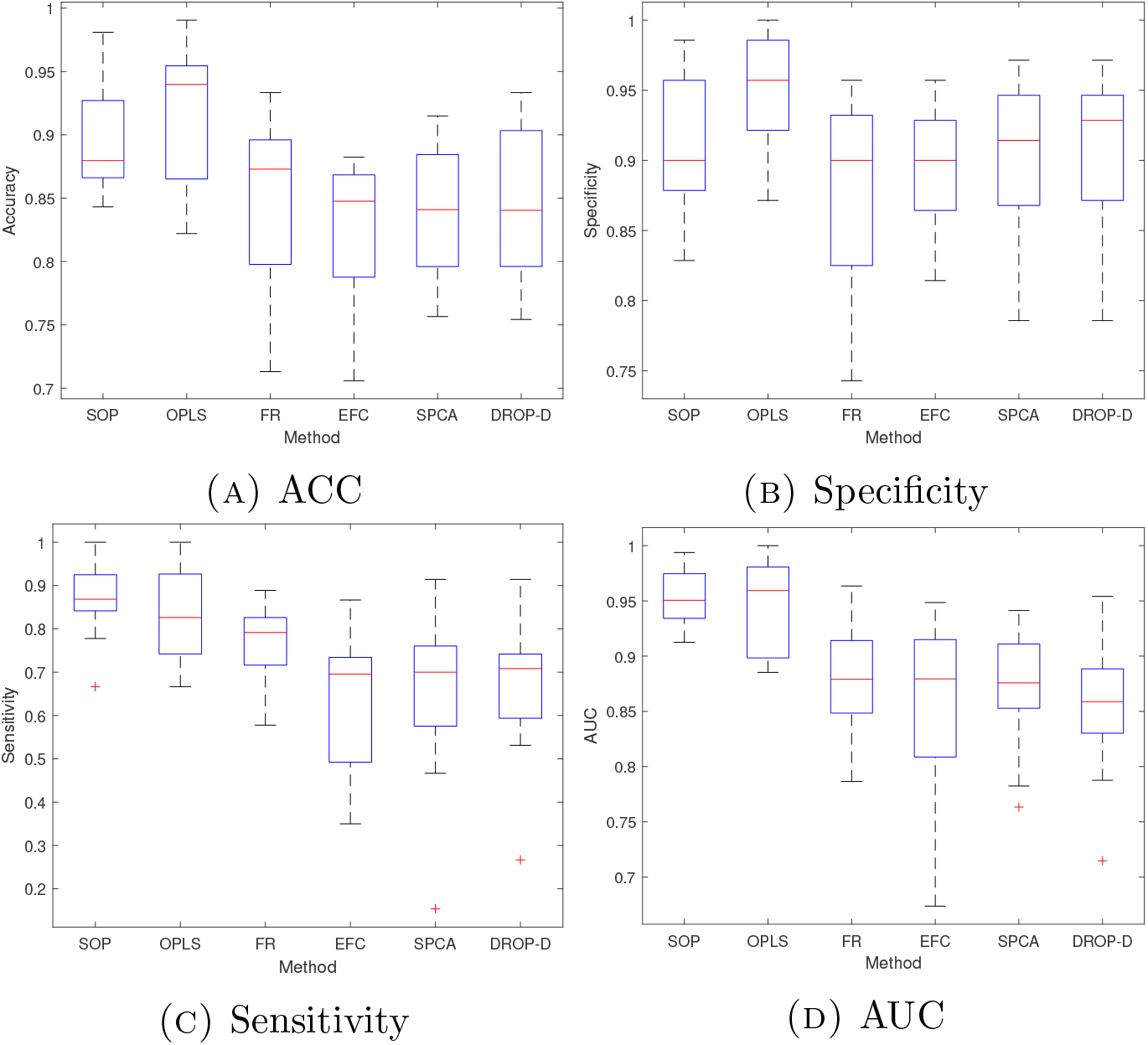
Box plots showing the median, upper and lower quartiles, and minima and maxima, across 13 classification results on the Keller data set. The outliers are highlighted as red crosses.

**Table 5.**
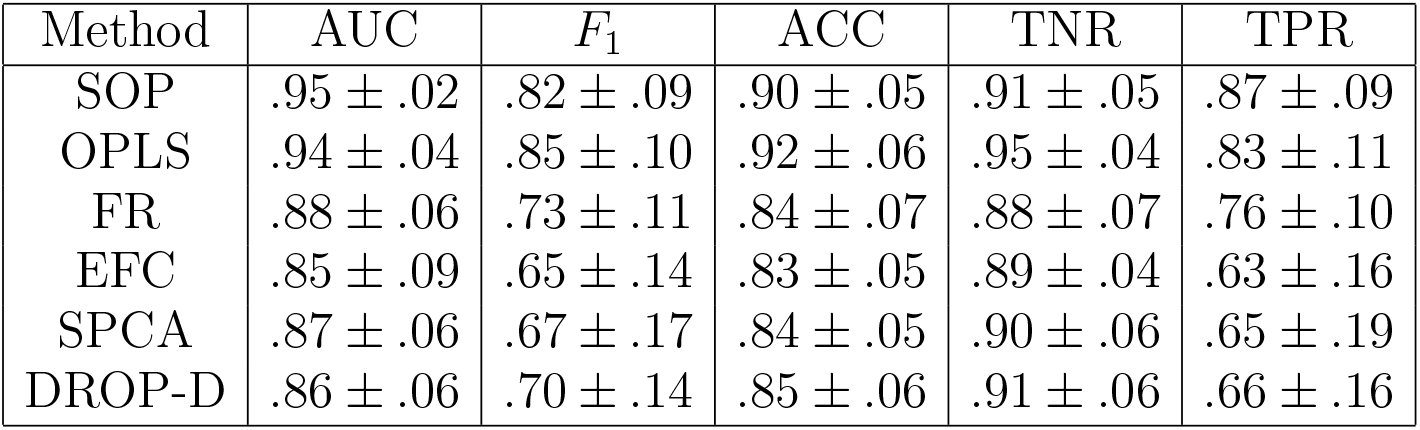
Keller *et. al*. results.

##### B.2.2. Results - Chan et. al. data

In this section, we present our results on the data provided by Chan *et. al*. in [2]. i.e., the data discussed in point (4) of section A.5. In figure 7, we present an ROC plot comparison using SOP, OPLS, FR, EFC, SPCA, and DROP-D. In table 6 we report the classification scores discussed in section A.3.

**Figure 7.**
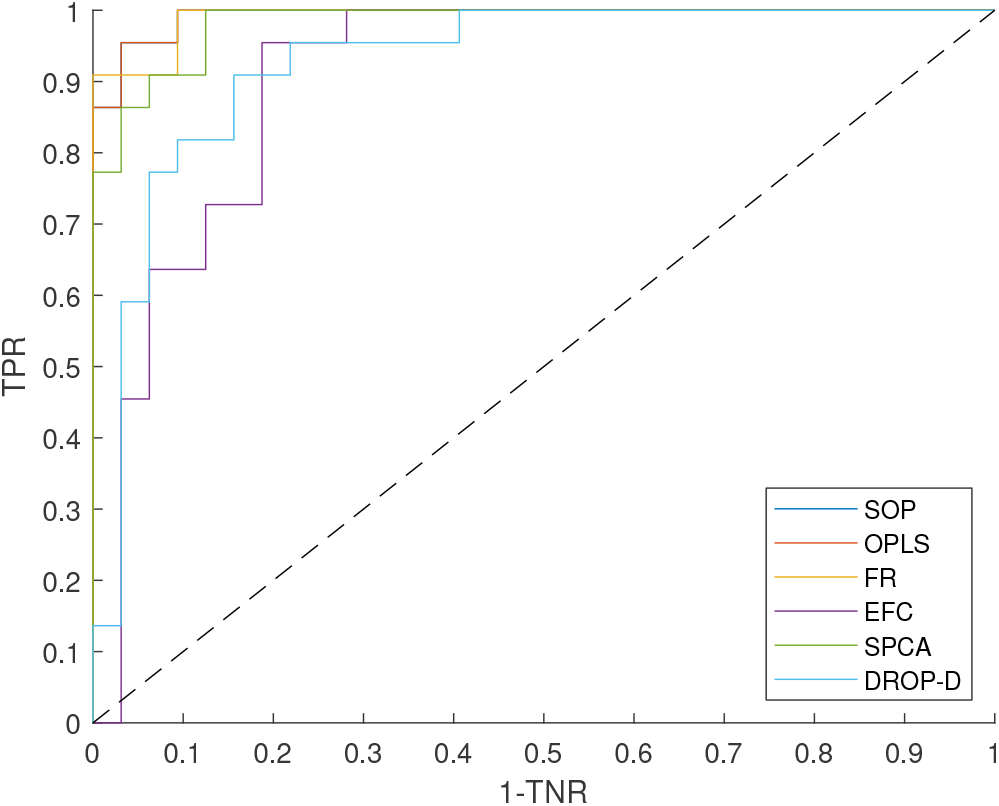
ROC plot comparison on the Chan *et. al*. data. The classification results for OPLS and SOP were identical in this case (see table 6), and hence the dark blue ROC curve (for SOP) is hidden behind the red ROC curve (for OPLS), as they match up exactly.

**Table 6.** Chan *et. al*. data results. In this table, 1 indicates that all 22 out of 22 healthy control patients were classified correctly. As we cannot reasonably claim a specificity of 1 on such a small sample size, we present the result as ∼ 1 to bring the readers attention to the small sample size.

| Method | AUC | $F_1$ | ACC | TNR | TPR |
| --- | --- | --- | --- | --- | --- |
| SOP | .99 | .95 | .94 | .95 | .94 |
| OPLS | .99 | .95 | .94 | .95 | .94 |
| FR | .99 | .95 | .94 | $\sim 1$ | .91 |
| EFC | .91 | .83 | .80 | .73 | .84 |
| SPCA | .98 | .92 | .91 | .86 | .94 |
| DROP-D | .93 | .89 | .87 | .82 | .91 |

In this example, the three methods which offer the best performance, are SOP, OPLS, and FR, with FR offering higher specificity at the cost of some sensitivity and vice versa for SOP and OPLS. The classes are more balanced here (32 cases and 22 controls) and the dimension *p* = 280 is significantly lower than the previous examples. Thus, the sparsity penalties are likely not as valuable here as in the case of the Lee *et. al*. data, for example, where the number of samples (*n* = 117) was significantly lower than the dimension (*p* = 2578). This example was included to show what happens for smaller *p*, and for more balanced class sizes. SOP offers a similar level of performance when compared the literature here, and thus SOP performs well and consistently in the case of smaller *p* and balanced class sizes.

#### B.3. Addressing the tuning of *k*

As noted in section A.9, the number of features (*k*), which yielded the best results in terms of the mean AUC score after cross validation, was chosen for the results presented throughout this section. The best *k* was chosen from *k* 2, …, 10. Although with such minimal tuning the level of overfitting is negligible, in this section we present our results using SOP when *k* is kept fixed at *k* = 3 to show what happens when a consistent model architecture for SOP is used throughout, and to be sure that there is no overfitting due to the choice of *k*. The remaining hyperparameters (such as the sparsity parameter *β*) are selected as discussed in section A.9. See table 7 where we have presented the results using SOP, with *k* = 3 fixed, on all data sets (1)-(4) discussed in section A.5. We see a very comparable performance to the best *k* values when *k* = 3 is fixed throughout, and SOP retains the performance advantages when compared to OPLS, FR, EFC, SPCA, and DROP-D. The tuning of *k* was done mainly to allow the feature selection methods, namely FR and EFC, a chance to perform well, as often more features than *k* = 3 are needed to achieve high performance. For example, in [19], a selection of miRNA panels are compared against with *k* ranging from *k* = 1 up to *k* = 10. In that paper, the larger *k* panels (e.g., *k* = 8) offered an improved performance when compared to the panels with smaller *k* (e.g., when *k* = 3).

**Table 7.** Results using SOP with *k* = 3 fixed throughout all classifications. The Japan and Keller *et. al*. data results are presented as in tables 1 and 5, and show the mean and standard deviation results across all classifications conducted in each case. The Lee *et. al*. and Chan *et. al*. data results are presented as in tables 3 and 6.

| Data set | AUC | $F_1$ | ACC | TNR | TPR |
| --- | --- | --- | --- | --- | --- |
| Japan | $98 \pm .03$ | $.88 \pm .12$ | $.99 \pm .01$ | $\sim 1 \pm .01$ | $.86 \pm .14$ |
| Keller *et. al.* | $.95 \pm .02$ | $.82 \pm .09$ | $.90 \pm .05$ | $.91 \pm .05$ | $.87 \pm .09$ |
| Lee *et. al.* | .98 | .96 | .94 | .86 | .97 |
| Chan *et. al.* | .99 | .94 | .93 | .91 | .94 |

## Appendix C

### Analysis of [19, 16, 14] data

In figure 8, we have presented scatter plots of the Japanese data using the *t*-distributed Stochastic Neighbor Embedding (TSNE) algorithm [17]. In figure 8a, the TSNE plot shows two distinct classes of controls, one high risk (mixed in with the cancer patients), and one low risk (some distance from the cancer patients). The low and high risk controls, the liver infections, and cancers are all highlighted by different colors in figure 8a, and labeled in the figure legend. The liver infections (e.g., cirrhosis and hepatitis) also formtheir own class far from the cancers and controls. We find that the control separation is largely due to hsa-miR-4783-3p, which has high correlation (*R* = .91) with the high risk/low risk control separation. In figure 8b we have plotted a histogram of the hsa-miR-4783-3p expression values across all control patients. We see a clear bimodal distribution, with one peak corresponding to low risk controls (the left-hand peak), and one to high risk controls (the right-hand peak). Such separation in the control set is not reported in [19, 16], nor is hsa-miR-4783-3p selected as a biomarker in the cancer predictions. It would seem that hsa-miR-4783-3p is worth investigating further. However, we choose to leave this for further work as the focus of this paper is testing performance of the new dimensionality reduction methodology.

**Figure 8.**
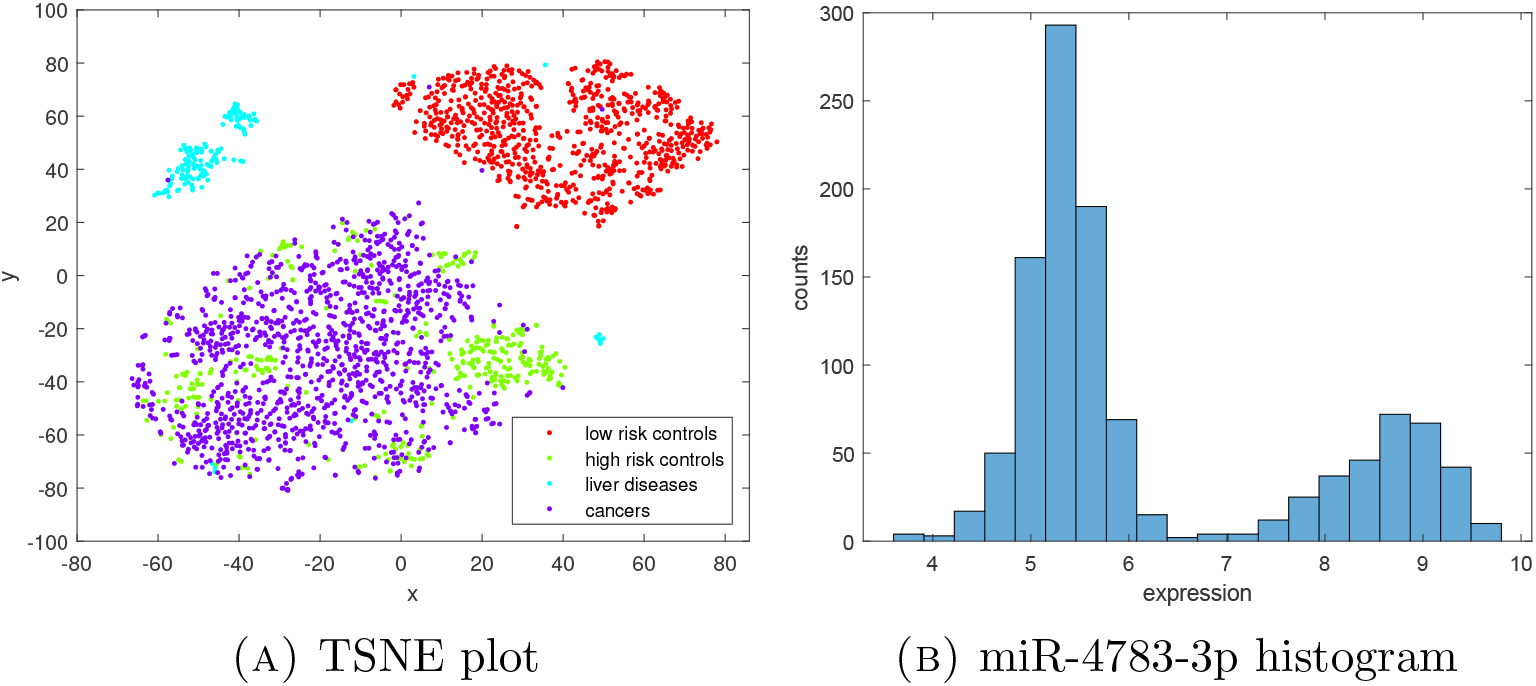
TSNE plot of Japan data and an expression histogram of hsa-miR-4783-3p, which is highly correlated (R = .91) to the separation between high risk and low risk controls. The left-hand mode of the miR-4783-3p histogram corresponds to the low risk controls, and the right-hand mode to the high risk controls.

#### C.1. Breast cancer patient clustering

Here we use TSNE to visualize the control samples, the cancer samples of [19, 16], and the advanced breast cancer cases of [14] in 2-D space. See figure 9. We notice that the cancer samples of [19, 16] form a different class to the advanced cancer samples of [14].

**Figure 9.**
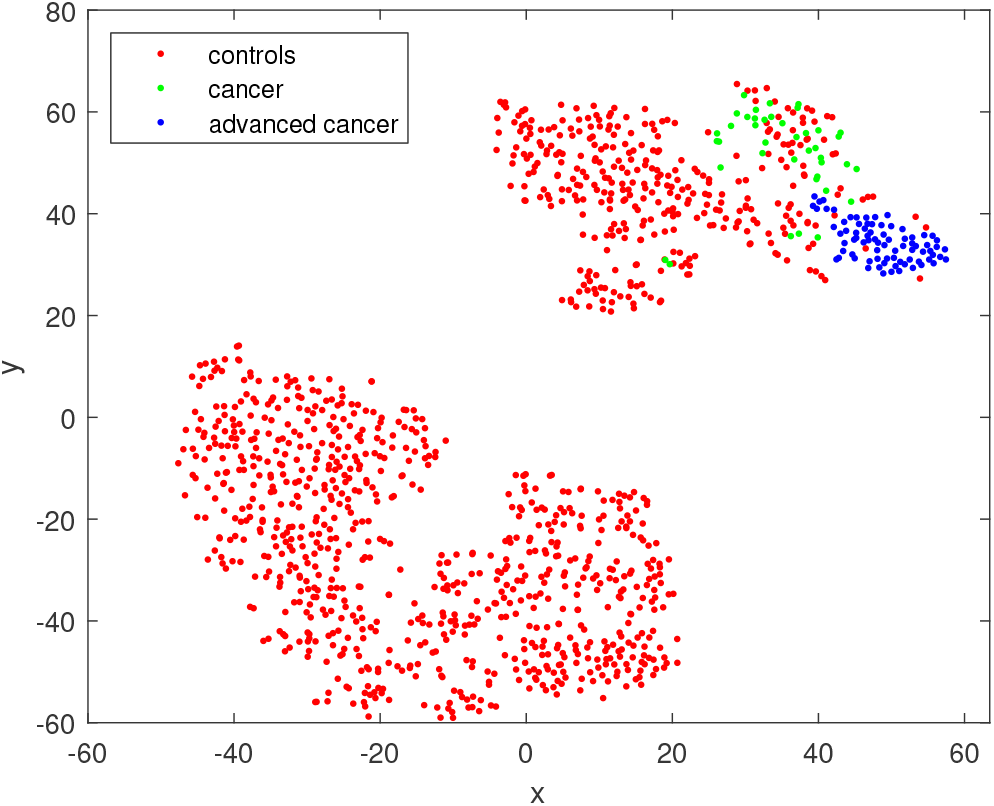
TSNE plot of Japanese control set and breast cancer samples. We have highlighted the breast cancer samples of [19, 16] in green, and the advanced breast cancer sample of [14] in blue.

## Notes

### Competing Interest Statement

The authors have declared no competing interest.

